# Long-term changes in the breeding biology of a New Zealand bellbird population suggest plasticity in life-history responses to ecological restoration

**DOI:** 10.1101/2021.01.18.427196

**Authors:** Michelle M. Roper, M.T. Harmer Aaron, Dianne H Brunton

## Abstract

Ecological restoration projects provide excellent opportunities to study how animals adapt their life-history strategies in response to changeable environments. A fundamental way animals can optimise reproductive success in changing conditions is trading-off aspects of their breeding system. The New Zealand bellbird (*Anthornis melanura*) has had a long-term presence on the small restoration island, Tiritiri Matangi Island (Tiri), spanning the island’s degraded agricultural past to its current extensively restored state. We studied the breeding biology of this bellbird population to assess how their reproductive life-history strategies have responded over time to the restoration on Tiri. We compared the current breeding data (2012–2016) of the bellbirds with data from between 2001–2010 (including Baillie, 2011, Cope, 2007), and from 1977–1978 (Anderson and Craig, 2003), prior to the island’s restoration. We also explored associations between abiotic/biotic factors and bellbird reproductive success for the most recent period (2012–2016). Our main finding was that clutch size significantly declined over time from a mean of 3.6 to 2.4 eggs per nest and this decline correlated with increasing population density. This is consistent with a density dependent effect, although further data are required to empirically test this conclusion. Overall, the earliest spring laying dates were in late August and the latest extended to January, with all chicks fledged by the end of February. Nest success was 47% (range 40 – 54%) across 2012–2016, falling within a similar range as previous studies. We found little effect of year, weather, parental age or morphometrics on reproductive success. We observed directional change in patterns of parental investment between 1977–1978 and 2012–2016; in 2012–2016, parents persisted with raising single broods rather than abandoning and re-nesting to raise larger broods. These results suggest that the bellbirds’ life-history traits are plastic in response to local conditions which provides an advantage when repopulating a regenerating or changing habitat.

## Introduction

Islands restored to their native habitat and a functioning ecological state are crucial for the long-term conservation of New Zealand’s endemic flora and fauna. New Zealand (hereafter NZ) has been a pioneer in island conservation since its mainland forest cover was reduced from 75% to as little as 25% following human settlement (Saunders and Norton, 2001). Monitoring animal and plant populations on these restoration islands is also essential to understand the viability of populations and their suitability as sources for future translocations. Valuable insights come from studies of native species protected on these restoration islands, especially studies on reproductive life-history of populations. For example, reproductive success is not density dependent in small island populations of North Island (NI) robin (*Petroica longipes*) and hihi (*Notiomystis cincta*) on the scientific reserve of Tiritiri Matangi Island (Armstrong and Ewen, 2013), but on Mokoia Island, a small population of NI saddleback (*Philesturnus carunculatus*) appears to be density dependent, with reduced reproductive output within 10 years of establishing on the island (Davidson, 1999). Although these results reflect how a species’ breeding biology can adjust with conspecific density, breeding biology may also be influenced by improved ecosystem function. Longer-term data for species on restoration islands are needed to assess how a population’s reproductive life-history traits can respond to a restoring ecosystem.

Species with life-history traits that are ‘phenologically flexible’, such as the ability to adjust reproductive timing in response to changing temperatures (Moussus et al., 2011), are likely to better adjust and persist in changing environments. Restoration islands are a special case of rapidly changing habitats, and success of reproductive trade-offs in life history strategies is likely to be strong in these populations, as the environment and population density can change rapidly over time. As the habitat restores, the availability of food and nest sites is expected to increase. Changes in the amount or timing of food availability will allow parents to invest energy into producing more or higher quality young over potentially longer breeding periods (Martin, 1987). However, as the density of a population increases, reproductive output may reduce due to density-dependence and increased competition for limited resources (Lack, 1954). Density-dependent reproductive output occurs as birds are either forced into lower quality habitat or are faced with an increase in competition that causes a uniform decrease in habitat quality (Both, 1998, Ferrer and Donazar, 1996). Competition due to higher densities can lead to an increase in nest failure (Arcese et al., 1992, Both, 1998), reduction in territory size, and/or an increase in parental foraging time (Sillett et al., 2004). To compensate for high levels of competition, reproductive trade-offs are made to ensure an individual’s reproductive output is maximised whilst minimising costs on their own survival (Martin, 1987, Sæther et al., 1996). For example, birds can first reduce egg volume to conserve energy but produce the same number of offspring (Martin, 1987). However, there is a limit to which the egg volume can reduce, as mortality of the young will increase, and thereby a reduction in clutch size may better optimise the birds’ reproductive output (Martin, 1987). These trade-offs can also be driven by environmental changes that include food availability, climate, and predation (Camfield et al., 2010, Martin, 1987, Martin, 1995, Sandercock et al., 2005).

Reproductive output is also optimised by an individual’s ability to use different strategies with respect to timing of breeding, breeding attempts per season, clutch size and parental investment. Breeding is ideally timed so that the periods of highest food demand by chicks coincide with abundant food availability in the environment (Tomás, 2015). To achieve this, birds can alter incubation timing and intervals (Crick et al., 1993, Tomás, 2015). For example, multi-brooders (ability to raise multiple successful broods) can start breeding earlier but they will have larger clutches with increasing food availability, then clutch size declines over the breeding season as food availability reduces (Crick et al., 1993). Low food availability can delay the start of breeding and reduce reproductive output (Marshall et al., 2002), whereas in superabundant food years, individuals can have earlier laying dates and an increase in clutch size (Hoi et al., 2004). Parents must also consider their condition and predation risk; hence, female clutch size can change to alter number of feeding visits to the nest (Lack, 1954) and parents can reduce nest visits when predation risk is high (Kleindorfer, 2007, Olsen et al., 2008). Reproductive output is hence linked to how flexible an individual’s response is to changing environments and conditions.

An individual’s reproductive success can also be influenced by other factors including phenotypic traits and short-term weather events. Phenotypic traits can have an effect on reproductive output; these include the parents’ body morphometrics (Baran and Adkins-Regan, 2014, Cain and Ketterson, 2012, Langston et al., 1990, McDonald et al., 2005) and age (Bédard and LaPointe, 1985, Forslund and Pärt, 1995, Imlay et al., 2017, Jankowiak and Wysocki, 2016, Jarvinen, 1991, Lack, 1954, Nol and James, 1987, Pärt, 1995). It has been suggested that selection should favour smaller female size as smaller females have lower energetic requirements and could hence breed earlier (Downhower, 1976), but in some cases, such as red-winged blackbirds (*Agelaius phoeniceus*), larger females lay earlier and have higher nest success (Langston et al., 1990). However, whether female size has a causal effect on reproductive success is debateable as size can correlate with other traits such as aggressiveness (Cain and Ketterson, 2012, Langston et al., 1990). Short-term weather effects, such as total rainfall over a year or less, can have both positive and negative effects on reproductive success. For example, food availability (Chamberlain et al., 1999) and nest survival (Chase et al., 2005, Fantle-Lepczyk et al., 2016) can depend on higher rainfall.

However, total rainfall during certain stages of the nesting period can be detrimental to nestling survival and fledging success (Arlettaz et al., 2010, Fantle-Lepczyk et al., 2016, Öberg et al., 2015). Higher temperatures prior to fledging can counteract the effect of higher rainfall early in the nesting period in Eurasian hoopoes, *Upupa epops*, with an increase in nestling survival (Arlettaz et al., 2010). Therefore, understanding how phenotypic traits and short-term weather variables influence reproductive success will also need to be considered when examining reproductive trade-offs in response to a changing environment.

We studied the plasticity of critical components of reproductive life history traits, particularly clutch size, for the New Zealand bellbird (*Anthornis melanura*; hereafter bellbird) on a restoring conservation island by using short-term and long-term data. The bellbird is a socially monogamous songbird in the family Meliphagidae (honeyeaters) that has benefited from restoration projects. The bellbird occurs throughout most of the islands of New Zealand but has undergone local extinctions, such as the northern third of the North Island by 1870 (Bartle and Sagar, 1987). The restoration of conservation managed islands has allowed bellbirds to flourish. For example, on Tiritiri Matangi Island the bellbird population declined to less than 24 birds by the time farming ended in 1972 (Baillie, 2011, Cameron and Davies, 2013), but then increased after intense habitat restoration, mammalian predator eradication and supplementary feeding (Baillie, 2011, Graham and Veitch, 2002, Roper, 2012). As the bellbird population has grown and other avian re-introductions have occurred, bellbirds may now be experiencing increasing competition for resources. Using data on bellbird reproduction from 1977 to 2016 to examine long-term changes in bellbird reproductive life history traits, we predicted clutch size would decline over time, possibly due to both intra- and inter-specific competition for resources. We also examined what factors, including environmental and parental morphometrics, could influence aspects of short-term reproductive success. We predicted that short-term variations in nesting success would be associated with rainfall; with higher rainfall leading to greater food abundance.

## Methods and materials

### Study site and species background

Our study was conducted on the scientific reserve Tiritiri Matangi Island (hereafter Tiri). Tiri is a low-lying 220 ha island in the Hauraki Gulf, 28 km north of Auckland, New Zealand (36.60° S, 174.89° E). Prior to the study in 2010, the island had a bellbird population of approximately 6.3–10 birds per ha (Roper, 2012). Bellbirds have been present on the island for at least 100 years but only in the last 30 years has the bellbird population expanded after intensive replanting and invasive mammal eradication (Rimmer, 2004). The island is now representative of a typical northern New Zealand coastal forest with ephemeral flowering and fruiting plant species providing food resources year-round (Gravatt, 1970).

Bellbirds are an endemic New Zealand honeyeater. They are sexually dimorphic in plumage (males are darker olive green and females have a white cheek stripe) and size, with females approximately 20% smaller (Heather and Robertson, 2005). Their song is sexually dimorphic with males tending to sing longer song bouts and females singing single song types with more variable intervals (Brunton and Li, 2006). They exhibit social monogamy, and although both sexes feed the chicks, females undertake more chick feeding and solely carryout incubation (Cope, 2007, Heather and Robertson, 2005). Their diet consists of nectar, seasonably available fruit, and invertebrates (Craig et al., 1981, Rasch and Craig, 1988, Roper, 2012). There are relatively few studies on their breeding biology (Anderson and Craig, 2003, Cope, 2007, Massaro et al., 2008) and no studies have compared changes in nesting biology over time.

### Data collection

We collected breeding data from 2012 to 2016 as part of a larger study on bellbird behavioural ecology that began in 2001. We compared these data with data collected from other published and unpublished studies (see Table 1). The bellbird breeding season occurs during the austral spring and summer, generally from September to February (Heather and Robertson, 2005). Nests were located by finding pairs on territories and observing the birds for nesting behaviours (e.g. carrying nesting material) and searching for built nests. We regularly checked nests (every 2–4 days) to record start and end dates for each nesting stage (building, laying, incubation and hatching) to determine nesting timing and variation between the years. We also measured clutch size, brood size and the number of fledged chicks to calculate hatching and fledging success. Nest site selection measures included nest height and nest plant species. We were limited to measuring nests at a maximum height of 8m (but note that most nests were below this limit).

**Table 1.** Nesting data for bellbirds on Tiritiri Matangi Island from current (2012–2016) and previous (1977–2010) studies.

| Year | Total<br>number of<br>nests | Clutch size |  |  | Brood size |  |  | Hatching<br>success |  | Fledging<br>success |  | Apparent<br>nest success |  | Source |
| --- | --- | --- | --- | --- | --- | --- | --- | --- | --- | --- | --- | --- | --- | --- |
|  |  | *N* | Mean | Median | *N* | Mean | Median | *N* | % | *N* | % | *N* | % |  |
| 1977/78 | 27 | 16 | 3.60 ± 0.10 |  | 16 | 2.56 ± NA |  | 16 | 75 | 16 | 68 |  |  | Anderson and Craig,<br>2003 |
| 2001 | 20 |  | 3.00 ± 0.10 |  | 18 | 2.40 ± 0.2 |  | 18 | 83 | 16 | 64 | 19 | 79 | Oron, T. and Brunton,<br>D.H. Unpublished |
| 2005/06 | 54 | 45 | 2.75 ± 0.08 | 3 |  |  |  |  |  |  |  |  |  | Cope, 2007 |
| 2005 | 33 |  |  |  | 20 | 1.50 ± 0.24 |  | 33 | 85 |  |  | 23 | 65 | Cope, 2007 |
| 2006 | 21 |  |  |  | 16 | 1.40 ± 0.29 |  | 21 | 62 |  |  | 20 | 55 | Cope, 2007 |
| 2007 | 20 | 11 | 2.91 ± 0.09 | 3 | 16 | 1.81 ± 0.31 | 2 | 11 | 69 | 10 | 85 | 14 | 64 | Baillie, 2011 |
| 2009 | 32 | 19 | 2.52 ± 0.12 |  | 19 | 1.20 ± 0.12 |  |  |  | 13 | 66 |  |  | Brunton, D.H.<br>Unpublished |
| 2010 | 15 | 15 | 2.44 ± 0.24 |  | 9 | 1.70 ± 0.24 |  |  |  | 9 | 71 |  |  | Brunton, D.H.<br>Unpublished |
| 2012 | 52 | 24 | 2.63 ± 0.15 | 3 | 17 | 2.35 ± 0.23 | 2 | 17 | 89 | 17 | 90 | 39 | 77 | This study |
| 2013 | 81 | 50 | 2.32 ± 0.10 | 2 | 47 | 2.21 ± 0.12 | 2 | 47 | 95 | 35 | 85 | 58 | 72 | This study |
| 2014 | 54 | 26 | 2.23 ± 0.16 | 2 | 25 | 2.00 ± 0.19 | 2 | 25 | 91 | 22 | 80 | 36 | 58 | This study |
| 2015 | 68 | 38 | 2.45 ± 0.11 | 3 | 37 | 2.24 ± 0.14 | 2 | 37 | 91 | 23 | 90 | 55 | 62 | This study |
| 2016* | 112 | 4 | 2.00 ± 0.41 | 2 | 4 | 2.00 ± 0.41 | 2 | 4 | 100 | 4 | 62.5 | 103 | 65 | This study |

\* Smaller sample size occurred for specific clutch and brood sizes due to most nests observed from a distance as we did not have experienced nest monitoring personnel at the site for long periods.

Where possible, we captured and banded the adults using mist nets or specially modified sugar-water feeder traps near the nests. The birds were banded with unique combinations of colour and metal bands and as part of our banding protocol we measured weight (g) with Pesola scales, tarsus length (mm) and head-bill length (mm) with Vernier calipers, and wing length (mm) and tail length (mm) with a wing rule. We also collected data on their age (hatch year (HY), second year (SY) or after second year (ASY), determined by plumage and the presence or absence of a wing slot; Craig, 1985). All capture, handling and banding protocols were conducted under permits from the New Zealand Department of Conservation (20666-FAU, 34833-FAU, 2008/33) and the Massey University Animal Ethics Committee (12/32, 15/21). We avoided capturing around the nests when we knew the female was incubating, but for females caught with an engorged brood patch, we banded and weighed the bird but bypassed other measurements to reduce handling time.

### Nest parameters and nest success

We compiled nesting data from 2012 to 2016 into one database. This included timing of nesting, breeding stage of nest when found and ended, nest site parameters (location, nest height, nest plant species), outcome of nest (success/failure), predation of nest, clutch size, number of unhatched eggs, brood size, number of fledglings, age of fledglings if known, and parent’s identification if banded, along with their body morphometrics. Weather data (daily rainfall and daily maximum and minimum temperatures - CliFlow; https://cliflo.niwa.co.nz/) were sourced from a weather station on the island monitored by the Department of Conservation. When data were not available on the island, we used data from the neighbouring Whangaparoa station on the mainland (approximately 5.65 km away). We obtained the bellbird density and population estimates for Tiri from the following sources for each year: 1977 (Anderson and Craig, 2003), 2003 (Kevin Parker, unpublished data), 2010 (Roper, 2012), and 2015–2016 from the Supporters of Tiritiri Matangi Island (John Stewart, unpublished data).

We investigated the relationship between various abiotic/biotic factors and clutch size, brood size and nest success. We tested for short-term differences in clutch size, brood size and plant species used for nest sites among years (2016 excluded for clutch and brood size due to low sample size) with Kruskal-Wallis tests. We also tested for a change in clutch size from 1977 to 2015 with a Kruskal-Wallis test. Regression models were fitted to investigate the relationships between clutch size and population size over time in the program Curve Expert Professional (Hyams, 2018). We chose the best regression models based on the smallest standard error (SE), highest r^2^ value and smallest 95% confidence interval range. We compared the plant species selected for nest sites between years with a chi-square test and nest height was compared between years with a one-way analysis of variance (ANOVA). We compared fledging success, as the mean ± SE number of chicks fledged, against clutch size and brood size with a Kruskal-Wallis test. For all statistical analyses in this study, we used a significance threshold of *P* = 0.05. To assess seasonal reproductive output per female, we calculated the mean ± SE number of eggs laid, and chicks fledged, per female (only banded females known to have multiple clutches).

To investigate the effects of the parental body morphometrics on clutch size and number of fledglings, we reduced the dimensionality of four morphometric measurements (wing, tail, head-bill and tarsus length) using a principle component analysis (PCA; Table 2). The PC scores were used in a Pearson correlation in R (R Core Team, 2015) with clutch size and number of fledglings. Parents that were measured outside of the breeding season and in moult were excluded from the PCA analysis. We then compared the parents’ body morphometrics (wing, tail, head-bill and tarsus length) between the sexes with Pearson correlations to investigate patterns of assortative mating. We used a Mann-Whitney U test to compare second year (SY) first-time breeders and after second year (ASY) adults for differences in start date of incubation, clutch size, brood size and number of fledged chicks.

**Table 2.** Loadings of the four morphometric measures on PC1 and PC2 from the PCA analysis for female and male bellbirds.

|  |  | PC1 | PC2 |
| --- | --- | --- | --- |
| Female | Wing | -0.532 | 0.485 |
|  | Tail | -0.574 | -0.040 |
|  | Tarsus | 0.207 | 0.868 |
|  | Head-bill | 0.587 | 0.095 |
| Male | Wing | -0.662 | 0.001 |
|  | Tail | -0.636 | 0.055 |
|  | Tarsus | 0.118 | 0.976 |
|  | Head-bill | 0.378 | -0.212 |

We calculated nest success, in terms of a nest successfully fledgling at least one chick, as apparent nest success (number of nests that fledged chicks divided by total number of nests) and a corrected version using daily survival rate (DSR) of nests. DSR was calculated as an adaptation of the maximum likelihood estimator for calculating nest success (Bart and Robson, 1982, Johnson, 1979), which was originally developed in the program MARK (Rotella, 2007, Rotella et al., 2004, Rotella, 2017). This method produces comparable results to other methods of calculating nest success, such as Stanley’s method, as these methods take into account exposure days which reduces bias from the observers missing nests that failed early on, unlike apparent nest success (Jehle et al., 2004). DSR of nests (period from first egg laid to fledging) was calculated using the R package RMark (Laake, 2013). We modelled DSR with year, total monthly rainfall, maximum temperature and minimum temperature to investigate what variables may best explain DSR. Nest success from start to finish was then calculated by raising the DSR to the power of the number of days from egg laying to fledging (Rotella, 2017). We compared levels of nest failure caused by predation during laying, incubation and fledgling period. Other causes of nest failure were combined as we often could not be certain of the cause (e.g. desertion and storm events). We also compared nest failure at different nest heights.

## Results

### Timing of breeding

A total of 367 nest attempts were observed over the five breeding seasons from 2012 to 2016. The earliest observed building attempt was recorded on 3 September 2013 (Figure 1), although one female was found already incubating on 2 September 2013. Where the start date of incubation was not known, it was estimated from the average incubation duration (from clutch completion to last egg hatching) of 12.4 ± 0.3 days (*N* = 14), with a range of 11 to 14 days. Length of nestling period (from all eggs hatched to all chicks fledged) was 14.4 ± 0.7 days (*N* = 17) and ranged from nine to 21 days (excluding nests that fledged early due to known disturbance). Initiation of incubation varied among years (Figure 1) with the earliest attempts beginning in early September in 2013 and 2015, which were years of abundant New Zealand flax (*Phormium tenax*) flowering (personal observation). There was no consistent pattern of peak nesting activity; peaks varied with year and in some year’s multiple peaks (e.g. 2014) were observed, while in others nesting activity was spread out across the entire breeding season (e.g. 2012; Figure 1).

**Figure 1.**
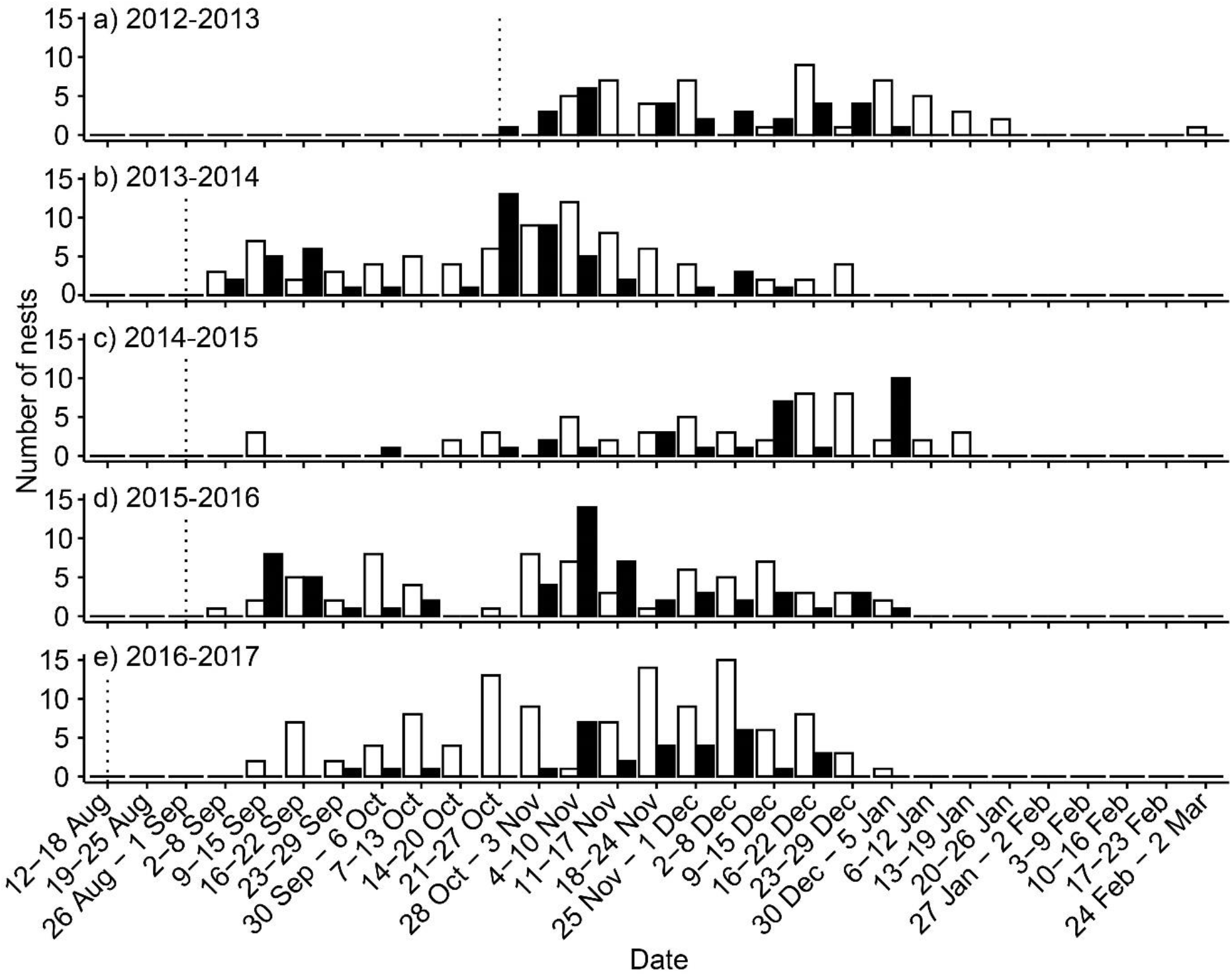
Number of nests found each week (unfilled bars) and number of nests starting incubation each week (filled bars) for each breeding season (a-e), with the week searching began (dotted line).

### Clutch size, brood size and number of fledglings

Among 142 nests where females laid eggs and where clutch size was known, median clutch size was 2, with a mean of 2.38 ± 0.06 and range of 1 to 4 eggs. There was no significant difference in clutch size among years 2012–2015 (Kruskal Wallis test, χ^2^ = 4.57, df = 3, *P* = 0.20). However, data from previous studies ranging back to 1977 suggests that clutch size has declined over time (Figure 2; Table 1). Comparing the raw data over all years except for 2001, 2009 and 2010 which could not be obtained, there was a significant decline in clutch size from 1977 to 2015 (Kruskal Wallis test, χ^2^ = 45.87, df = 9, *P* < 0.001). Population size on the island grew exponentially over this period from 1977 to 2015 (r^2^ = 0.77, SE = 422.5), whereas clutch size reduced following a Gaussian function (r^2^ = 0.91, SE = 0.14; Figure 3). Hatching success ranged from 89–95% (excluding 2016 due to small *N*) with previous studies ranging from 64–83% (Table 1).

**Figure 2.**
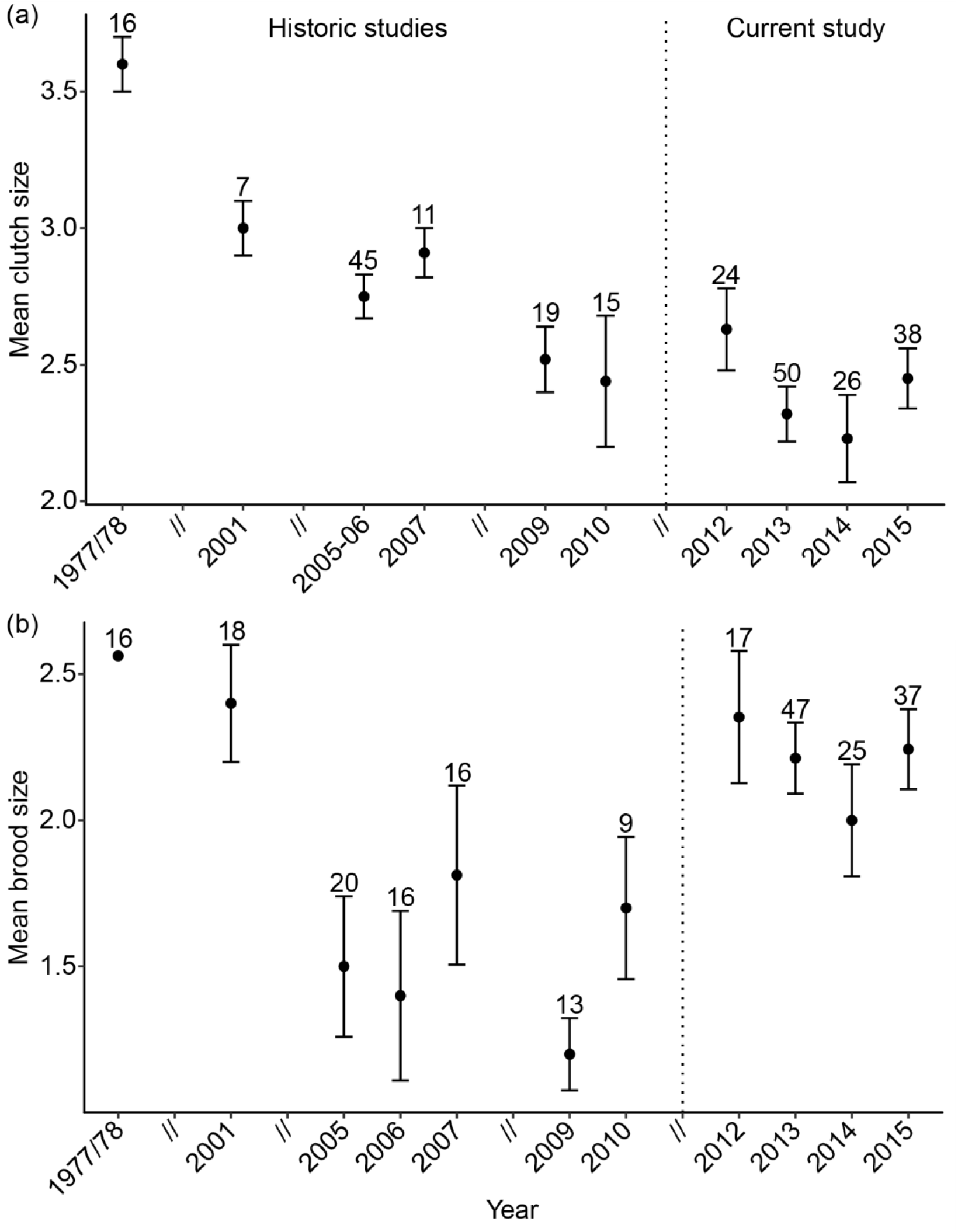
Mean clutch size ± SE (a) and mean brood size ± SE (b) with *N* values for bellbirds from current study and studies going back to 1977. See Table 1 for references.

**Figure 3.**
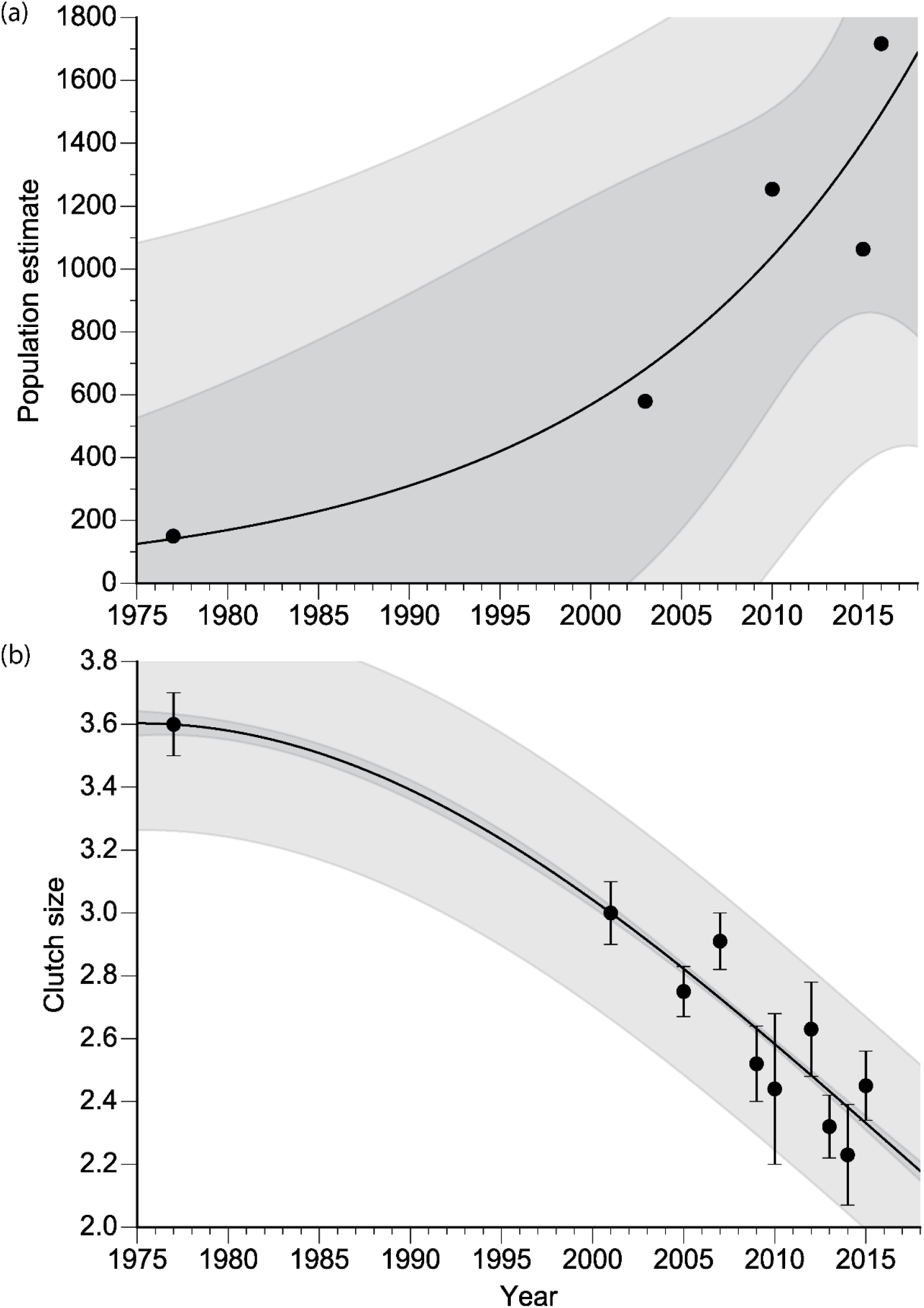
Non-linear regression models for (a) the Tiri population estimate (exponential) and (b) clutch size (Gaussian model) over time with the 95% confidence interval (dark grey) and prediction interval (95% confidence; light grey).

For our data set, 130 nesting attempts had at least one egg hatch successfully. Median brood size was 2, with a mean of 2.19 ± 0.08 and range of 0 to 4 chicks. Mean brood size for our data set was relatively high compared to studies from 2005 to 2010, but similar to earlier studies (Figure 2). There was no significant difference in brood size among years 2012 to 2015 (Kruskal Wallis test, χ^2^ = 2.23, df = 3, *P* = 0.52; Figure 2; Table 1). The mean number of fledglings per active nest was 1.39 ± 0.08 (median 2, range 0 to 4) with also no significant difference among years (Kruskal Wallis test, χ^2^ = 3.83, df = 3, *P* = 0.28). Fledging success ranged from 80–90% (excluding 2016 due to small *N*) with previous studies varying from 47–92% (Table 1). As clutch size increased for individual females, the rate at which chicks fledged the nest significantly increased (Kruskal Wallis test, χ^2^ = 6.56, df = 2, *P* = 0.038) but not for brood size (Kruskal Wallis test, χ^2^ = 2.81, df = 2, *P* = 0.25). Across multiple broods (banded females only), the mean total number of eggs per female per season was 3.83 ± 0.53 eggs (*N* = 12) and mean total number of fledglings per female per season was 2.45 ± 0.41 (*N* = 20).

### Parental associations with breeding parameters

Comparisons of nesting parameters with adult body morphometric data did not show any strong correlations. Correlations of PC1 and PC2 with clutch size and number of fledglings showed only a weak positive correlation between female size (represented by PC1) and clutch size (Figure 4). There was no significant difference between nesting parameters of second year (SY) first time breeders and adults over 2 years old (ASY) for either sex, however ASY birds did tend to start nesting earlier, had larger clutch and brood sizes and more chicks fledged (Table 3). There was no correlation between the body morphometrics of males and females within breeding pairs (wing r = −0.09, *P* = 0.62; tail r = 0.12, *P* = 0.51, head-bill r = 0.28, *P* = 0.11; tarsus r = −0.21, *P* = 0.23).

**Table 3.** Comparison of reproductive performance between second year (SY; first breeding season) and after second year (ASY) for female and male bellbirds with a Mann-Whitney U test.

|  |  | SY |  | ASY |  | U | P |
| --- | --- | --- | --- | --- | --- | --- | --- |
| | | N | Mean ± SE | N | Mean ± SE | | |
| Female | Start of incubation (weeks)* | 10 | 11.70 ± 1.13 | 64 | 10.03 ± 0.10 | 260.5 | 0.35 |
| | Clutch size | 7 | 2.00 ± 0.22 | 53 | 2.38 ± 0.10 | 245.5 | 0.13 |
| | Brood size | 7 | 2.00 ± 0.22 | 51 | 2.12 ± 0.13 | 201.5 | 0.57 |
| | Number fledged | 10 | 1.00 ± 0.37 | 73 | 1.34 ± 0.13 | 427.5 | 0.37 |
| Male | Start of incubation (weeks)* | 8 | 11.25 ± 1.98 | 67 | 10.32 ± 0.53 | 247.5 | 0.73 |
| | Clutch size | 7 | 2.00 ± 0.31 | 57 | 2.49 ± 0.09 | 268 | 0.11 |
| | Brood size | 7 | 2.00 ± 0.31 | 57 | 2.23 ± 0.12 | 231.5 | 0.41 |
| | Number fledged | 4 | 1.00 ± 0.40 | 81 | 1.62 ± 0.12 | 215 | 0.23 |

\*Number of weeks from first nest found incubating.

**Figure 4.**
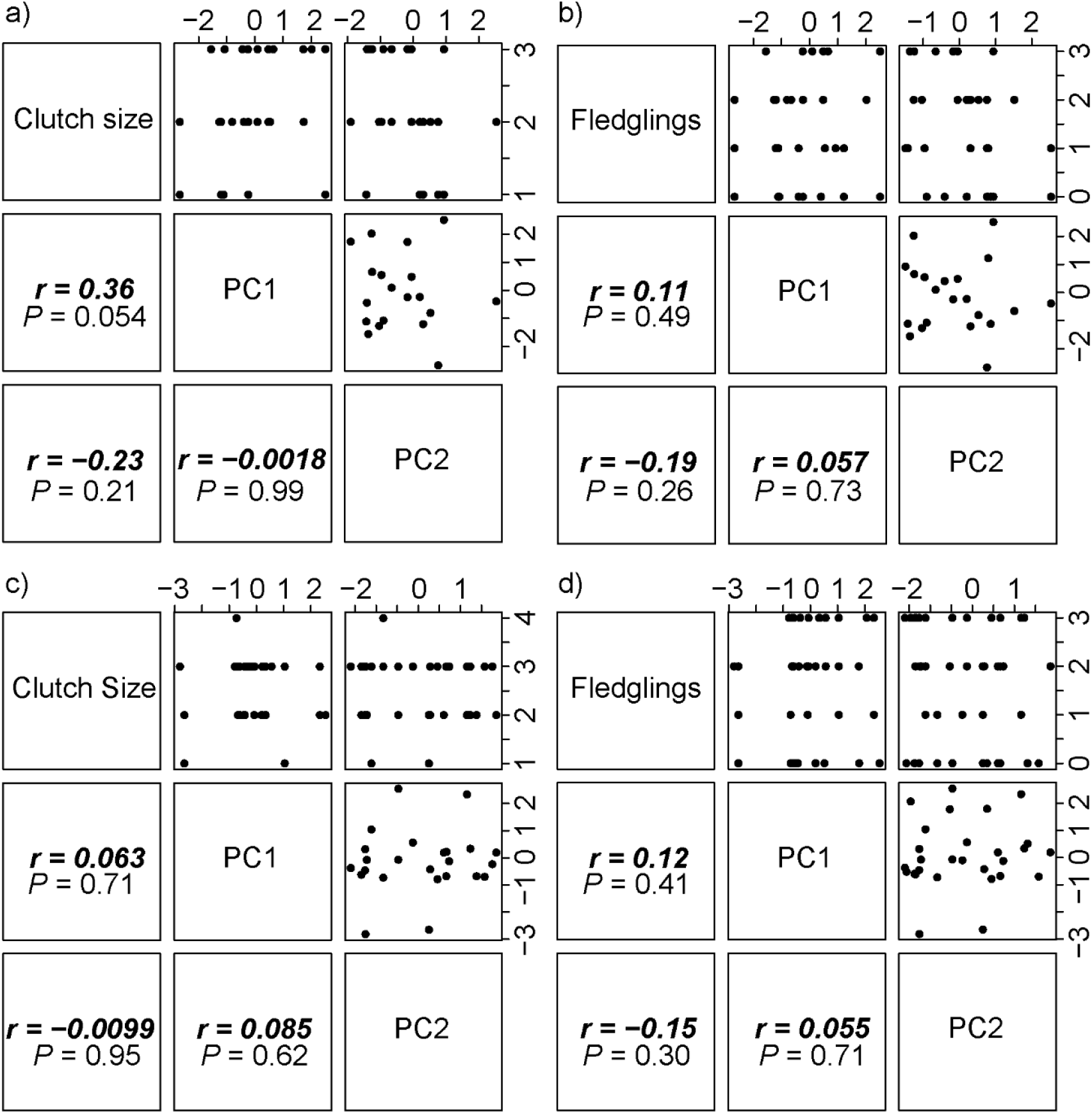
Correlation matrix for PCA analysis of morphological parameters, clutch size and number of fledglings for female (a, b) and male (c, d) bellbirds. Table 2 indicates the loadings of the four morphometric measures on PC1 and PC2.

### Nest site selection

Across 2012–2016, we found there was a significant difference between the proportions of each plant species used for nesting each year (χ^2^ = 1608.60, *df* = 76, *P* < 0.001). We most commonly found nests in the vines of *Muehlenbeckia complexa* and *M. australis* (range of 19.8 – 42.3% across 2012 to 2016; see supplementary Table S1 for full results). *Muehlenbeckia* often grows on other species, and we found nests most often with this genus in trees such as *Cordyline australis* (9.1 – 50.0%), *Pittosporum crassifolium* (0.0 – 36.4%) and *Metrosideros excelsa* (0.0 – 36.4%; Supplementary Table S1). Nest height ranged from 0.3 m to 8.0 m with a mean of 4.08 ± 0.13 (*N*=185). Nest height did not significantly vary between years (ANOVA, *F*(4) = 0.0023, *P* > 0.05).

### Nest success

Apparent nest success was relatively high, ranging from 58 to 77% and was similar to previous studies which ranged from 55 to 79% (Table 1). Daily nest survival rate (DSR) was around 97% for all years (Table 4). When all years were modelled together, year and weather parameters had lower rankings than constant daily survival rate (DSR ~ 1) and did not have a significant effect on DSR (Table 4). Based on average duration of laying, incubation and nestling periods, we calculated that a nest must survive 28 days to successfully fledge young. Nest success (i.e. proportion of nests fledging at least one chick) varied from 40% to 54% among years and was 47% when modelled across 2012–2015 (Table 4). This fell within the range for previous studies from 41–53% (Table 4).

**Table 4.** Daily nest survival rate (DSR) with model outcomes, and nest success for historic studies (1977–2006) and current study (2012–2015 plus all years combined).

| Breeding season | *N* | DSR estimate<br>± SE | Nest<br>success<br>(%) | Method | Source |
| --- | --- | --- | --- | --- | --- |
| 1977–78 | 17 | NA | 44 | Stanley’s | Anderson and<br>Craig, 2013 |
| 2001 | 19 | NA | 53 | Stanley’s | Oron, T. and<br>Brunton, D.H.<br>Unpublished |
| 2005 | 55 | NA | 45 | Stanley’s | Cope, 2007 |
| 2006 | 57 | NA | 41 | Stanley’s | Cope, 2007 |
| 2012 | 19 | 0.973 ± 0.011 | 48 | MARK |  |
| 2013 | 40 | 0.978 ± 0.005 | 54 | MARK |  |
| 2014 | 28 | 0.973 ± 0.007 | 47 | MARK |  |
| 2015 | 45 | 0.968 ± 0.006 | 40 | MARK |  |
| 2012–2015 | 132 | 0.973 ± 0.003 | 47 | MARK |  |
| Model | *k* | AICc | ΔAICc | *wi* | Deviance |
| DSR ~ 1 | 1 | 332.42 | 0.00 | 0.36 | 330.42 |
| DSR ~ year | 2 | 333.58 | 1.16 | 0.20 | 329.58 |
| DSR ~ minimum<br>temperature | 2 | 333.95 | 1.53 | 0.17 | 329.94 |
| DSR ~ maximum<br>temperature | 2 | 334.17 | 1.75 | 0.15 | 330.17 |
| DSR ~ total monthly<br>rainfall | 2 | 334.42 | 2.00 | 0.13 | 330.42 |

We found that nest predation occurred more frequently during the nestling stage compared to laying and incubation (Table 5). Over 80% of nest failures were caused by ‘other’ factors (such as desertion, storm events) and often could not be determined. For all other causes of nest failure, there was no consistent relationship across years between nest height and nest failure (Table 5).

**Table 5.**
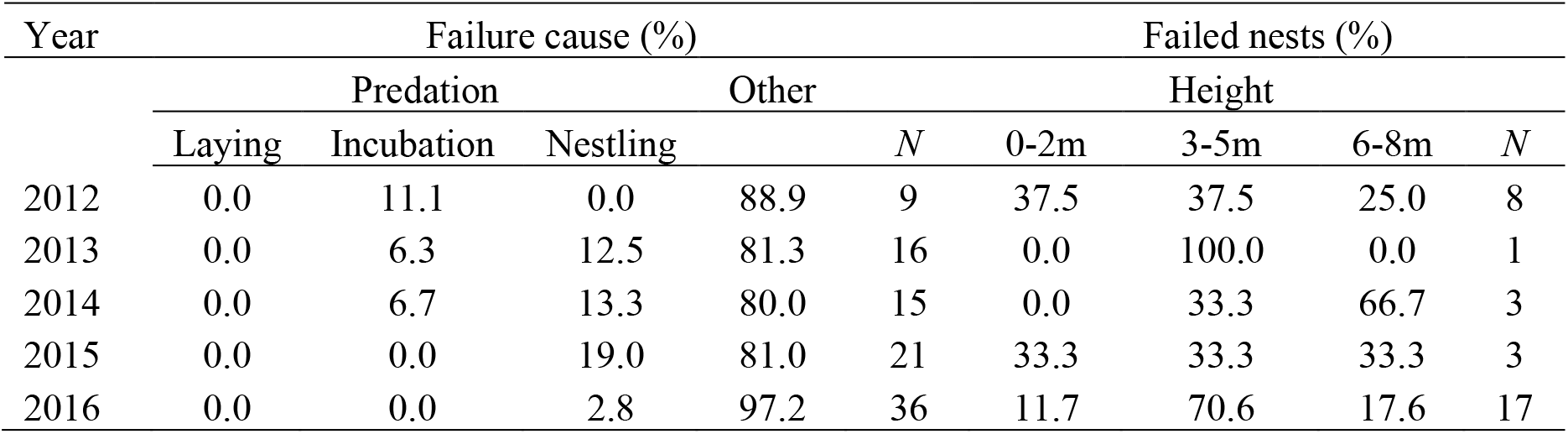
Causes of nest failure (proportion of nests) and proportion (%) of nests that failed at different nest heights.

## Discussion

In this study, we examined the current and historical breeding biology of a New Zealand bellbird population, located on an ecological restoration island, to investigate plasticity in reproductive life-history strategies. The traits that showed plasticity were 1) clutch size, evidenced by a long-term decline in clutch size, and 2) timing of breeding, evidenced by different temporal patterns of nesting between years. Plant hosts for nests varied in the short-term for recent years and differed from 1977-78 where tree ferns (e.g. *Cyathea medullaris*) were the most common nest site (Anderson and Craig, 2003). This variation in nest sites is likely a response to the regeneration of the flora on the island. Brood size appeared to fluctuate over time, but was overall higher in this study period, perhaps due to higher hatching success. Surprisingly, there was no apparent change in nest success (both apparent and adjusted) over time and no short-term variables had a significant effect on nest success and DSR. We are therefore uncertain why there was relatively higher hatching and fledgling success over our study period compared to previous years. A possible explanation for these higher values is that we had to exclude nests from the calculations where we were unable to determine if a nest failed during incubation, chick rearing or fledgling stages. Parental body morphometrics and age had no significant relationship with nesting parameters. However, female body size (PC1 for head-bill, wing and tail) had a weak positive association with clutch size. As no data were collected between 1978 and 2001, we cannot know if any parameters such as clutch size and nest success initially increased during the early phase of the restoration on Tiri. However, the results suggest that bellbirds were able to adapt to the changing conditions on Tiri. Importantly, current smaller clutch sizes suggest there may now be reproductive trade-offs limiting their reproductive output.

### Flexibility in bellbird breeding ecology

The mean clutch size of bellbirds on Tiri from 2012 to 2016 was the lowest on record since 1977 (Anderson and Craig, 2003). The bellbird population on Tiri has increased significantly since the eradication of mammalian predators/competitors (e.g. kiore, *Rattus exulans*) and re-vegetation of the island with native flora (Graham and Veitch, 2002). Since 2000, year-round supplemental sugar-water has been available to the bellbirds as part of the recovery plan for hihi, *Notiomystis cincta* (Castro et al., 2003, Chauvenet et al., 2012), a nectar-feeder slightly larger than the bellbird (Castro et al., 2003, Chauvenet et al., 2012). The bellbird population has grown from about 150 individuals in 1977/78 (Anderson and Craig, 2003) to an estimated 1200–2000 individuals in 2010 (Roper, 2012). The bellbirds on Tiri have likely benefitted from the provision of sugar-water as they use it intensively in the non-breeding season (Roper, 2012), which may have reduced winter mortality (as evidenced by hihi mortality reducing during the experimental provisioning of sugar-water on Tiri; Armstrong and Ewen, 2001). The population has grown exponentially and this large bellbird population is now likely under *K*-selection (Saether et al., 2016b); where density-dependent mortality of juveniles and adults may be regulating population growth (Saether et al., 2016a, Saether et al., 2016b). We suggest that this has led to an increase in competition for resources during the breeding season (Evans et al., 2005) and female bellbirds have reduced clutch size as a trade-off for optimal fitness (Arcese et al., 1992, Haccou and McNamara, 1998, Murphy, 2000, Pettifor et al., 2001).

The Tiri population appears to have become density-dependent at a lower population density than a more northern population. Aorangi Island (part of the Poor Knights Islands) is approximately 122 km north of Tiri and has a higher bellbird population density at 11 to 18.1 birds per ha (Sagar and Scofield, 2006) versus 7.6 birds per ha on Tiri (John Stewart, personal communication). (Although the Aorangi bellbird population is recognised as a subspecies, *Anthornis m. obscura*, recent studies suggest there is insufficient evidence to support this; Baillie, 2011.) Yet, Tiri appears to have become density-dependent at a lower density as shown by the trade-off in clutch size, which is lower than on Aorangi (2.87; Sagar, 1985). A key difference between these two islands is in the number of broods produced each season; Tiri bellbirds produce multiple broods compared to single broods on Aorangi (Sagar, 1985). The result is a generally higher annual reproductive output by bellbirds on Tiri. Another difference is that the Tiri population typically has a longer breeding season than the Aorangi population (Sagar, 1985). More favourable breeding conditions on Tiri over a longer breeding season may mean Tiri females can afford to trade-off clutch size but have multiple broods; whereas Aorangi females have larger clutches to compensate for only having time for one successful brood in a shorter season with favourable breeding conditions (Sagar, 1985). This could allow these two populations to optimise their reproductive output while using two different life-history strategies to cope with different environments.

The bellbirds’ flexibility in life history traits, such as reproductive trade-offs, is a significant advantage in changeable environments. Islands are vulnerable to changing conditions from biotic and abiotic factors. The bellbirds on Tiri appear to be able to adjust the timing of breeding as an additional trade-off to deal with these variable conditions. They were able to have extended breeding seasons that started as early as late August and finish in February. However, in some years such as 2014 and 2016, the breeding season had a delayed start and similar timing as found by Anderson and Craig (2003) in 1977–78. The longest breeding seasons (2013 and 2015) coincided with abundant NZ flax flowering years (personal observation), a common food source, but further testing is needed to assess this link. Other biotic factors influencing reproductive trade-offs include presence of competitors, particularly other bird species. Tiri has a high diversity of bird species that are likely a source of competition for food resources, particularly invertebrates. There is also likely competition for other resources (e.g. nest sites), from the closely related tui (*Prosthemadera novaeseelandiae*), which is larger than the bellbird and behaviourally dominant (Rasch and Craig, 1988). Tui are rare visitors to Aorangi Island (McCallum, 1981), suggesting this population may have reduced inter-specific competition compared to the Tiri population. Abiotic factors that may affect reproductive trade-offs include short-term drought and long-term availability of water. While our DSR models did not show rainfall to be an important factor for nestling survival, it may be important for other factors such as invertebrate abundance (Chamberlain et al., 1999). There are multiple environmental factors that need to be tested to assess why bellbirds make reproductive trade-offs but results of this study consistently show that bellbirds have the flexibility to adapt to local conditions across New Zealand’s variable environments.

### Bellbird nest success

Various factors can affect nest success, but bellbirds appear to have had relatively consistent nest success over time. When kiore were present on Tiri, nest success was 44%, but since the eradication of kiore, there has not been a marked increase in nest success for bellbirds or other species on the island, such as North Island robins (Anderson and Craig, 2003, Armstrong et al., 2000). This suggests kiore were not a major nest predator or competitor for food resources. In all studies of bellbird breeding biology on Tiri, nest success has remained below 60%. The Swamp harrier (*Circus approximans*) and morepork (*Ninox novaeseelandiae*) were the most common predators present over this study (personal observation), but we were only able to account for up to 20% of nest failures due to predation. Anderson and Craig (2003) suggested that the marginal habitat on Tiri results in greater exposure of nests to storm events. We also found nests tended to fail after storm events, but we could not separate desertion from other causes of loss. Perhaps the regenerating plant growth that covers 64% of the island (Cameron and Davies, 2013) is still more vulnerable to storm damage (particularly on the exterior of the forest edges) than older forest growth.

We were unable to detect any strong relationships between the body morphometrics or age of parents and reproductive success. Female bellbirds only showed a weak positive trend between body size and clutch size. Clutch size can indicate female condition and hence those that can raise larger clutches have higher fitness (Pettifor et al., 2001). However, this weak trend for female bellbirds suggests other factors have more influence on reproductive success, as has been found for the positive relationship between female bellbird song complexity and fledging success (Brunton et al., 2016). For male bellbirds, we did not find any correlations between body size and reproductive output unlike in other species (Lemon et al., 1992, Møller, 1990). This could perhaps be due to high levels of extra-pair paternity in bellbirds (Cope, 2007), hence clutch size and fledgling success per nest may not reflect a male’s true reproductive output. Age did not have a consistent effect on breeding success, but it has been shown for several species that young birds tend to breed later (Bédard and LaPointe, 1985, Conrad and Robertson, 1993, Jankowiak and Wysocki, 2016, Jarvinen, 1991). While older bellbirds in our study tended to have higher reproductive output, the relationship was not strong. More research is needed, including annual reproductive success, extra-pair paternity and larger sample sizes, as success with age could be caused by differences in either competence (Nol and James, 1987, Pärt, 1995) or experience (Jankowiak and Wysocki, 2016).

Whilst we did not find strong evidence for factors that could influence nest success, we observed changes in parental investment over time. Anderson and Craig (2003) found that bellbirds abandon nests when there is only one chick remaining and never raise single-brood nests. This is likely a result of gaining better reproductive investment by re-nesting than by raising a single chick (Anderson and Craig, 2003). However, we observed 30 nests fledging single chicks. This suggests that conditions have changed in favour of raising one chick over re-nesting, as the costs of egg laying and incubation may be more expensive compared to raising a chick to independence (Monaghan and Nager, 1997). We speculate that this may be due to greater competition for resources, particularly invertebrates, which makes producing eggs for re-nesting more costly than raising a single chick. The increasing number of bird species re-introduced to the island (Galbraith and Cooper, 2013) could potentially be increasing inter-specific competition for invertebrates during the breeding season.

### Implications for species’ management

Bellbird breeding plasticity and success on Tiri shows how important restoring islands can be in re-establishing populations to the higher densities seen on islands that remain relatively untouched by human settlement. Similarly, high bellbird densities have been achieved in ‘mainland islands’ with predator proof fences and where introduced predators are reduced to very low levels, e.g. Tawharanui Regional Park (Auckland, New Zealand). However, density may not be completely indicative of reproductive success (Vickery et al., 1992) as there is relatively high genetic connectivity between bellbird populations (Baillie, 2011) and some sites could potentially act as sink populations. However, species such as the hihi, for example, have not yet reached a self-sustaining population on Tiri, despite having similar life-history traits, as they rely on supplemental feeding (Armstrong and Ewen, 2001, Castro et al., 2003). Hihi do differ in their reproductive strategies, being cavity nesters (Castro and Robertson, 1997), and also rely on artificial nest boxes on Tiri (Armstrong and Ewen, 2013). The laying date of hihi on Tiri has been found to not coincide with the optimal period for laying, showing they may not have adaptive and phenotypically plastic reproductive traits that suit conditions on Tiri (de Villemereuil et al., 2019). This suggests that hihi life history is less adaptive to early-regenerating habitat, while bellbirds on the other hand perhaps show more plastic life-history traits that allow their success in adapting to regenerating habitat. Monitoring reproductive success is hence a valuable tool in managing species’ survival and knowing how flexible their life-history traits are will help identify at what stage of an ecological restoration a population will have the best chance of becoming self-sustaining.

## Author contributions

Michelle M. Roper contributed to the design of the study and development of the methodology, conducted the fieldwork, managed field assistants, performed statistical analyses and wrote the manuscript. Aaron M. T. Harmer assisted with the analyses and writing the manuscript. Dianne H. Brunton designed the study, developed methodology, assisted with fieldwork, assisted with analyses and contributed to the writing of the manuscript.

## Acknowledgements

We thank B. Maste, K. Schor, X. Sleeuenhoek, C. Jense, N. Villing, J. Soethout (University of Applied Science in Agriculture, Food Technology, Environmental and Animal Sciences, Leeuwarden), and A. Lachaume (Montpellier SupAgro, Montpellier) for their contribution to collecting data from 2013 to 2016, and their respective institutions for facilitating their stay in New Zealand. We thank previous researchers who provided their data for this study, such as Talya Oron, Taneal Cope and Shauna Baillie. We also thank C. Smith, D. Smith, M. McCready, R. Shepherd, V. Franks, B. Evans and the many Tiri volunteers for their assistance in the field. We’d also like to thank Kevin Parker and John Stewart for providing data on the Tiri bellbird population size. Logistical support was provided by the School of Natural and Computational Sciences of Massey University, the Department of Conservation, and the Supporters of Tiritiri Matangi Island; we thank them for their invaluable support.

