## Supplementary material Table S1 for "Long-term changes in the breeding biology of a New Zealand bellbird population suggest plasticity in life-history responses to ecological restoration"

**Table S1.** Primary and secondary (host species for *Muehlenbeckia sp.*) plant species used for nest site selection across each breeding season.

| Plant species | 2012 | | 2013 | | 2014 | | 2015 | | 2016 | |
|---|---|---|---|---|---|---|---|---|---|---|
| Primary | *N* | % | *N* | % | *N* | % | *N* | % | *N* | % |
| *Muehlenbeckia australis/complexa* | 22 | **42.3** | 16 | 19.8 | 20 | **37.0** | 19 | **27.5** | 31 | **27.7** |
| *Cordyline australis* | 9 | 17.3 | 34 | **42.0** | 10 | 18.5 | 7 | 10.1 | 24 | 21.4 |
| *Kunzea ericoides* | 5 | 9.6 | 6 | 7.4 | 1 | 1.9 | 5 | 7.2 | 8 | 7.1 |
| Unknown | 5 | 9.6 | 0 | 0.0 | 0 | 0.0 | 4 | 5.8 | 20 | 17.9 |
| *Melicytus ramiflorus* | 3 | 5.8 | 6 | 7.4 | 7 | 13.0 | 5 | 7.2 | 10 | 8.9 |
| *Metrosideros excelsa* | 3 | 5.8 | 3 | 3.7 | 13 | 24.1 | 15 | 21.7 | 12 | 10.7 |
| *Knightia excelsa* | 2 | 3.8 | 3 | 3.7 | 1 | 1.9 | 3 | 4.3 | 1 | 0.9 |
| *Phormium tenax* | 1 | 1.9 | 6 | 7.4 | 1 | 1.9 | 3 | 4.3 | 1 | 0.9 |
| *Leptospermum scoparium* | 0 | 0.0 | 2 | 2.5 | 0 | 0.0 | 0 | 0.0 | 0 | 0.0 |
| *Myrsine australis* | 0 | 0.0 | 1 | 1.2 | 1 | 1.9 | 3 | 4.3 | 0 | 0.0 |
| Dead tree | 0 | 0.0 | 1 | 1.2 | 0 | 0.0 | 2 | 2.9 | 0 | 0.0 |
| Other (<2% each) | 2 | 3.8 | 3 | 3.7 | 0 | 0.0 | 3 | 4.3 | 5 | 4.5 |
| Secondary (*Muehlenbeckia sp.* host) | | | | | | | | | | |
| *Pittosporum crassifolium* | 8 | **36.4** | 1 | 7.1 | 0 | 0.0 | 2 | 14.3 | 0 | 0.0 |
| *Metrosideros excelsa* | 4 | 18.2 | 3 | 21.4 | 4 | **36.4** | 3 | 21.4 | 0 | 0.0 |
| *Cordyline australis* | 2 | 9.1 | 7 | **50.0** | 3 | 27.3 | 5 | **35.7** | 1 | 25 |
| *Kunzea ericoides* | 2 | 9.1 | 0 | 0.0 | 2 | 18.2 | 1 | 7.1 | 0 | 0.0 |
| *Coprosma sp.* | 2 | 9.1 | 0 | 0.0 | 1 | 9.1 | 0 | 0.0 | 2 | **50.0** |
| *Paraserianthes lophantha* | 2 | 9.1 | 0 | 0.0 | 0 | 0.0 | 0 | 0.0 | 0 | 0.0 |
| *Melicytus ramiflorus* | 1 | 4.5 | 1 | 7.1 | 2 | 18.2 | 2 | 14.3 | 0 | 0.0 |
| Dead tree | 1 | 4.5 | 1 | 7.1 | 0 | 0.0 | 1 | 7.1 | 1 | 25.0 |
| Unknown | 0 | 0.0 | 1 | 7.1 | 0 | 0.0 | 0 | 0.0 | 0 | 0.0 |
| *Vitex lucens* | 0 | 0.0 | 0 | 0.0 | 1 | 9.1 | 0 | 0.0 | 0 | 0.0 |

Note: Most commonly used species highlighted in bold for each breeding season.
